# High-sensitivity pattern discovery in large, paired multi-omic datasets

**DOI:** 10.1101/2021.11.11.468183

**Authors:** Andrew R. Ghazi, Kathleen Sucipto, Gholamali Rahnavard, Eric A. Franzosa, Lauren J. McIver, Jason Lloyd-Price, Emma Schwager, George Weingart, Yo Sup Moon, Xochitl C. Morgan, Levi Waldron, Curtis Huttenhower

## Abstract

Modern biological screens yield enormous numbers of measurements, and identifying and interpreting statistically significant associations among features is essential. Here, we present a novel hierarchical framework, HAllA (Hierarchical All-against-All association testing), for structured association discovery between paired high-dimensional datasets. HAllA efficiently integrates hierarchical hypothesis testing with false discovery rate correction to reveal significant linear and non-linear block-wise relationships among continuous and/or categorical data. We optimized and evaluated HAllA using heterogeneous synthetic datasets of known association structure, where HAllA outperformed all-against-all and other block testing approaches across a range of common similarity measures. We then applied HAllA to a series of real-world multi-omics datasets, revealing new associations between gene expression and host immune activity, the microbiome and host transcriptome, metabolomic profiling, and human health phenotypes. An open-source implementation of HAllA is freely available at http://huttenhower.sph.harvard.edu/halla along with documentation, demo datasets, and a user group.

**Author Summary:** Modern scientific datasets increasingly include multiple measurements of many complementary data types. Here, we present HAllA, a method and implementation that overcomes the statistical challenges presented by data of this type by using feature similarity within each dataset to find statistically significant groups of features between them. We applied HAllA to simulated and real datasets, showing that HAllA outperformed existing procedures and identified compelling biological relationships. HAllA is widely applicable to diverse data structures and presents the user with grouped results that are easier to interpret than traditional methods.

## Introduction

Pattern discovery in high-dimensional, heterogeneous data is a longstanding problem in applied statistics [1,2]. It is challenging for several reasons, including the inherent tradeoffs between sensitivity and generality - that is, the ability and power to detect associations given varying assumptions about the functional form of the relationship [3]. When applied in contexts such as high-throughput biology, these challenges are exacerbated by noisy, diverse, and non-linear data. Examples include biospecimens drawn from large cohorts, in which each sample may be decorated with heterogeneous phenotypic variables (clinical features, diseases status, etc.) and multiple high-dimensional molecular measurements (microbial taxa, epigenetic markers, gene expression, etc.). In the biological sciences specifically, selecting a subset of associations for follow-up validation experiments can be a complex yet important decision point. A gap remains to efficiently identify related features in such data, while both maintaining sensitivity and controlling spurious association reporting.

All-against-all (AllA) approaches, which test all pairs of features and then correct for false discovery, scale well only in completely independent tests of moderate size [4]. Under other conditions, such feature-wise approaches can have limited statistical power due to testing many correlated hypotheses for individually weak associations [5]. This has led to the development of a variety of (typically parametric) block-testing approaches, such as partial least squares (PLS) [6], canonical correlation analysis (CCA) [7], PLS discriminant analysis (PLS-DA), sparse principal component analysis (SPCA) [8], and SPARSE-CCA [9]. These serve to detect associations between reduced-dimensional representations of large input datasets, but they are typically limited by one or more of 1) applicability only to continuous measurements with no missing values (or only categorical, not mixed; PLS, CCA, SPCA); 2) a focus on the single, strongest axis of covariation between the datasets (CCA); 3) an assumption of linear covariation (CCA, SPCA, PLS); 4) identifying complex combinations of feature loadings implicated in associations, rather than specific features (particularly in kernel methods such as Kernel PCA [10]); and 5) a lack of explicit control of the false discovery rate (FDR).

Recent advances have focused on nonparametric methods for identifying highly general (i.e., linear and non-linear) associations between individual pairs of features, sometimes relying on computational or permutation-based methods not readily accessible to early applied statisticians. These include, for example, distance correlation (dCor) [11], which measures (not necessarily linear) dependency of two random variables with possibly different dimensions. The Chatterjee rank correlation (XICOR) [12] is another recently-introduced similarity measure that uses rank differences to assess the degree to which one variable is a measurable function of another. While dCor and XICOR provide comparatively general methods to discover complex associations between individual pairs of features, when applied to many combinations of linear feature pairs with varying degrees of dependence, the resulting statistical power can fall below simpler traditional approaches after controlling FDR for multiple hypothesis tests [13].

In this work, we develop a hierarchical all-against-all association testing framework (HAllA) that identifies highly general association types in paired, high-dimensional, and potentially heterogeneous datasets. HAllA preserves statistical power in the presence of collinearity by testing coherent clusters of variables in a hierarchical manner, while controlling overall FDR with hierarchical multiple hypothesis testing. HAllA discovers associations between blocks of features among paired datasets in a way that increases interpretability by grouping features according to their relatedness.

## Methods

In this section, we provide an overview of the HAllA algorithm and its component steps. Additional methods details, including pseudocode, are provided in S1 Appendix.

### The HAllA Algorithm

Hierarchical All-against-All Association testing (HAllA) identifies block associations between two potentially heterogeneous datasets co-indexed along one axis (Fig 1A). This co-indexing is referred to as the “samples” axis (columns), and the measurement axis as “features” (rows). For a pair of datasets containing measurements that describe the same set of samples and a specified pairwise similarity measure, the HAllA algorithm proceeds by 1) optionally discretizing features to a uniform representation (if required by the similarity measure), 2) finding the Benjamini–Hochberg (BH) FDR threshold, 3) hierarchically clustering each dataset separately to generate two data hierarchies, 4) coupling clusters of equivalent resolution between the two data hierarchies, 5) testing coupled clusters for statistically significant association in block format where the block passes a threshold for false negative tolerance (FNT), and 6) iteratively increasing resolution by descending through the pair of hierarchies according to which split results in a higher Gini score gain. The final pair of hierarchies are those that lead to the largest hypothesis blocks that pass the FNT threshold (Fig 1 and S1 Appendix).

**Figure 1.**
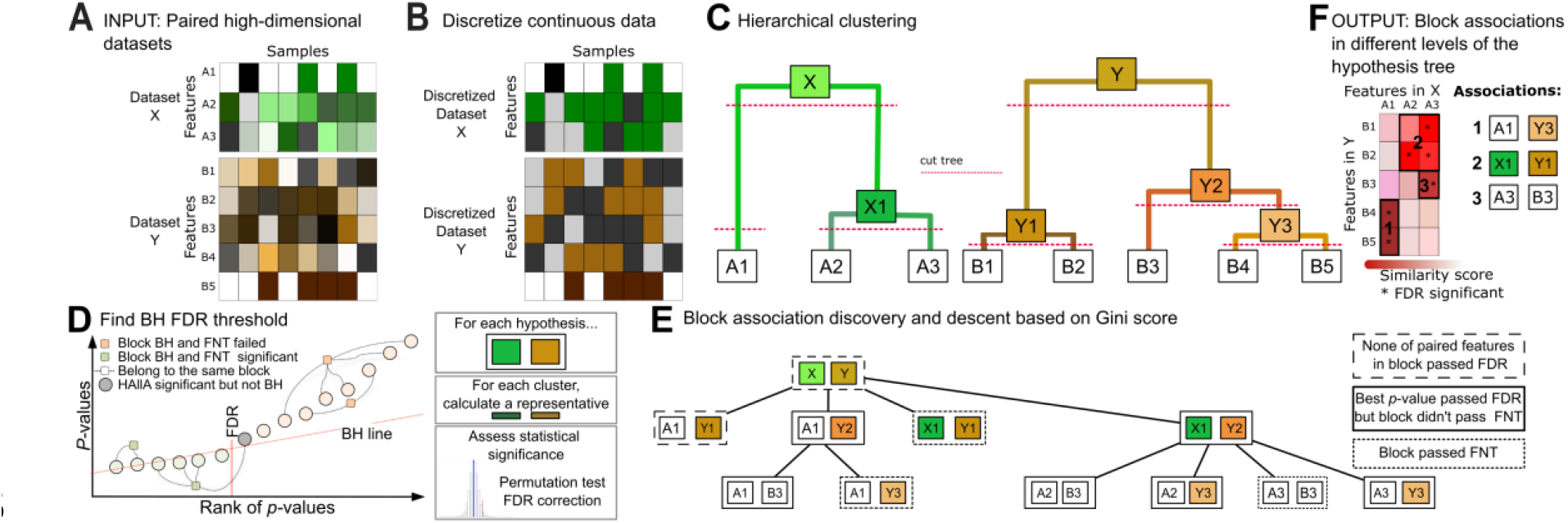
Hierarchical all-against-all (HAllA) association testing. **(A)** HAllA provides a novel method for heterogeneous association discovery in high dimensional data. Input data are represented in matrix form as features (rows) and samples (columns). **(B)** Data are discretized to provide a unified representation of heterogeneous feature types. This step is skipped for similarity metrics that requires continuous data (e.g. Spearman). **(C)** Features within each data set are hierarchically clustered using average linkage and Spearman association as default methods. **(D)** Reject block-wise null hypotheses that pass the false negative tolerance (FNT) threshold using Benjamini-Hochberg FDR threshold for pair-wise associations within the block. **(E)** Block format hypotheses are built by pairing clusters between two datasets at equivalent relative homogeneity. Each hypothesis node has two data clusters whose descendants are used for the next level of hypothesis testing. In hypothesis testing, the FNT threshold is used to determine which clusters are significantly associated between the two datasets. **(F)** Significant associations are reported after controlling the FDR for each hypothesis set in the descending approach using hypothesis tree-oriented structure.

### Optionally discretizing input datasets

This step permits direct comparison of continuous and categorical features (Fig 1B) and further enables the application of highly general measures of association from information theory, such as mutual information (MI). This combination allows HAllA to detect significant 1) non-linear associations between paired continuous features (e.g., quadratic or sinusoidal relationships), 2) differences in group means for paired continuous and categorical features, and 3) non-random associations between paired categorical features. HAllA’s default discretization scheme divides continuous features into bins of equal size once at the start of processing. By default, the number of bins is the cube root of the sample size, which provides reasonable power at a variety of sample sizes and correlation levels (Fig 1 in S1 Appendix). HAllA also removes features with low variance by applying a configurable frequency threshold (defaulting to 100%, meaning only features with no variability are removed) in order to reduce the number of unnecessary tests.

### Hierarchical clustering and cluster coupling allow detection of associations between groups of features

Each dataset is subjected to average-linkage hierarchical clustering using the specified similarity measure (Spearman’s rank correlation by default) within each dataset (Fig 1C). Associations between datasets are tested in a top-down manner by pairing nodes of similar resolution between their respective data trees. More specifically, HAllA recursively builds a tree of hypotheses to test (the “hypothesis tree”), beginning at the top of each dataset’s tree, descending to a set of nodes within each data tree, and then pairing each selected node from the first tree with each selected node of the second tree. At each step in the descent process, the choice of whether to descend within the X or Y hypothesis tree is made by comparing which split leads to a higher Gini score gain. In the case of ties, both descent steps are made. This procedure is repeated until termination, i.e. when the hypothesis block passes the FNT threshold or when the selected nodes represent single features in their respective data trees (Fig 1E). Another way to visualize this process is by focusing on the all-by-all hypothesis matrix (Fig 1F, left). The process begins by checking if the entire matrix passes the FNT threshold. If not, the matrix is recursively cut horizontally or vertically into smaller hypothesis blocks, with the position of each cut decided by each dataset’s similarity tree and Gini score gain. The cutting process stops when the smaller hypothesis blocks pass the FNT threshold or have been reduced to one-by-one blocks.

The notion of identifying and testing hypotheses in a hierarchical manner was previously proposed by Yekutieli [14]. HAllA’s hypothesis tree similarly groups more specific child hypotheses below a more general parent hypothesis. However, HAllA’s approach differs fundamentally from the Yekutieli approach in that HAllA tests hierarchical hypotheses until a null hypothesis can be rejected; Yekutieli’s method tests until the first failure to reject a null hypothesis. This results in HAllA maintaining greater power, while Yekutieli’s method instead maintains greater specificity.

### Determining the statistical significance of block associations

The method proceeds by testing the nodes in the hypothesis tree (each representing a pair of feature clusters, one from each dataset) for significant between-cluster associations. Each node in the hypothesis tree is evaluated using the following procedure: let ℋ denote the null hypothesis that the two clusters of features are not related, and ℋ_*i*_ be the null hypothesis of no association between two individual features within those clusters. Define *R*^*i*^ as the p-value of the association between an individual pair of features considered by ℋ_*i*_. We then count all rejected ℋ_*i*_ (i.e. *R*^*i*^ *≤ k*_*BH*_), and all ℋ_*i*_ that failed to reject, i.e. *R*^*i*^ *> k*_*BH*_ where *k*_*BH*_ is the global BH FDR threshold. The blockwise FNT is provided by the user (default FNT = 0.2) and acts as the allowed fraction of paired associations which are expected to fail to reject despite being true associations. If the fraction of paired associations in a block with *R*^*i*^ > *k*_*BH*_ is greater than or equal to FNT, we reject the entire block hypothesis ℋ.

If any hypothesis involved clusters rather than feature tips, and failed to reject, the procedure is repeated with new null hypotheses for associations between sub-clusters (Fig 1E), as described in section “Descending in sub-hypotheses of block hypotheses” in S1 Appendix. HAllA reports all significant associations between clusters of any size that pass the FNT threshold (Fig 1F).

### Visualizing outputs

Once the analysis is complete, the results are visualized in a “HAllAgram” (Fig 4). This comprises a heatmap visualizing the relatedness and strength of association between pairs of features in the two datasets. Features are ordered along each axis according to their position in the hierarchical tree so that clusters of significant features can be boxed into contiguous units. Marginally associated pairs are dotted, and each hypothesis block is labelled with the rank of its association strength. Features not associated with any block are not plotted by default. For analysis results where large numbers of blocks are detected, only the strongest blocks are shown (30 by default), with potentially-incomplete, lower-ranked blocks boxed in grey. Together, this set of plotting techniques allows users to visually understand the related sets of hypotheses that HAllA has detected. Other plotting utilities are also included with the method’s current implementation, such as a clustermap that displays the entire association tree in the margins for both datasets, as well as a diagnostic plot that displays the input data associated with individual hypothesis blocks.

## Results

### HAllA increases power while controlling FDR to report blockwise associations

When applied to paired datasets with no significantly related blocks of features, HAllA’s descent algorithm reduces to all-against-all (AllA) direct pairwise feature testing. In such circumstances, HAllA is expected to perform similarly to AllA. However, when there are sets of correlated variables within one dataset that are correlated with another set of variables in the other, HAllA will report the block-wise associations. Notably, we expect this behavior to be common in multi-omics data, where we see large clusters of molecular features (e.g. co-expressed genes in a metabolic pathway).

To evaluate these claims, we applied HAllA and AllA to paired, synthetic datasets generated with the data simulator function in the HAllA software. These datasets contained pre-specified block associations, which allowed us to evaluate the statistical and computational performance of these two methods (Fig 2 and Fig 3). With a constant target FNT in associated blocks of 0.2, HAllA controls FDR, reports association in block form, and improves power on average by 7-11% (Fig 2A) across varied FDR thresholds. HAllA also consistently boosts the true positive rate relative to AllA using different target FNT values in associated blocks (Fig 2B).

**Figure 2.**
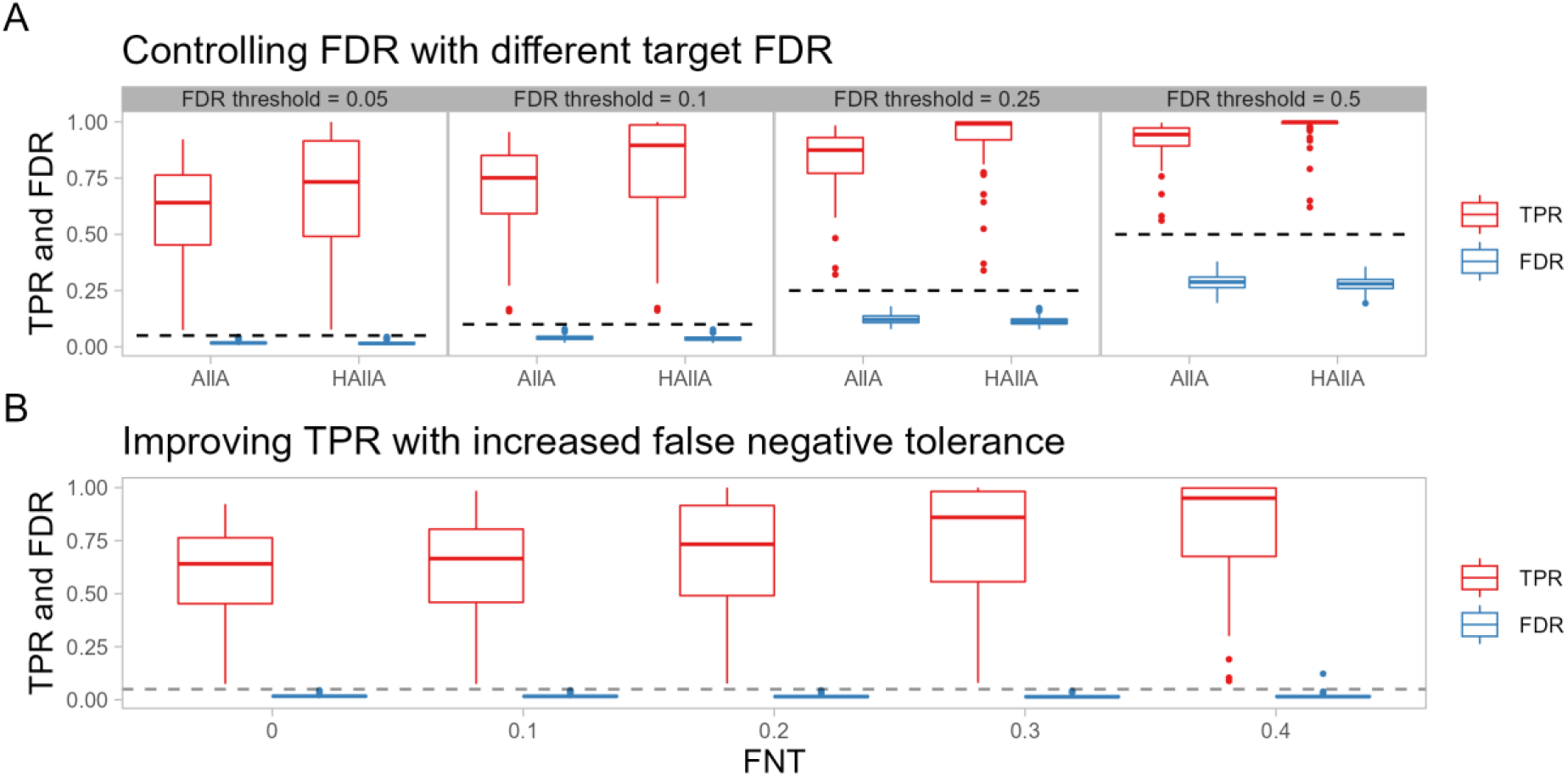
HAllA improves statistical power while controlling the FDR. 50 paired, synthetic datasets with 200 features and 50 samples containing clusters with linear block associations were analyzed. A) with FNT = 0.2, HAllA maintains the simulated FDR below the target (here (0.05, 0.1, 0.25, and 0.5), with associated trade-offs in statistical power. In addition, HAllA is consistently better powered than all-against-all (AllA) association testing across this range of target FDR values. Dashed lines parallel to the x-axis indicate the target FDR value in each comparison. B) By increasing the FNT, HAllA can improve the true positive rate with a comparatively minor increase in FDR.

**Figure 3.**
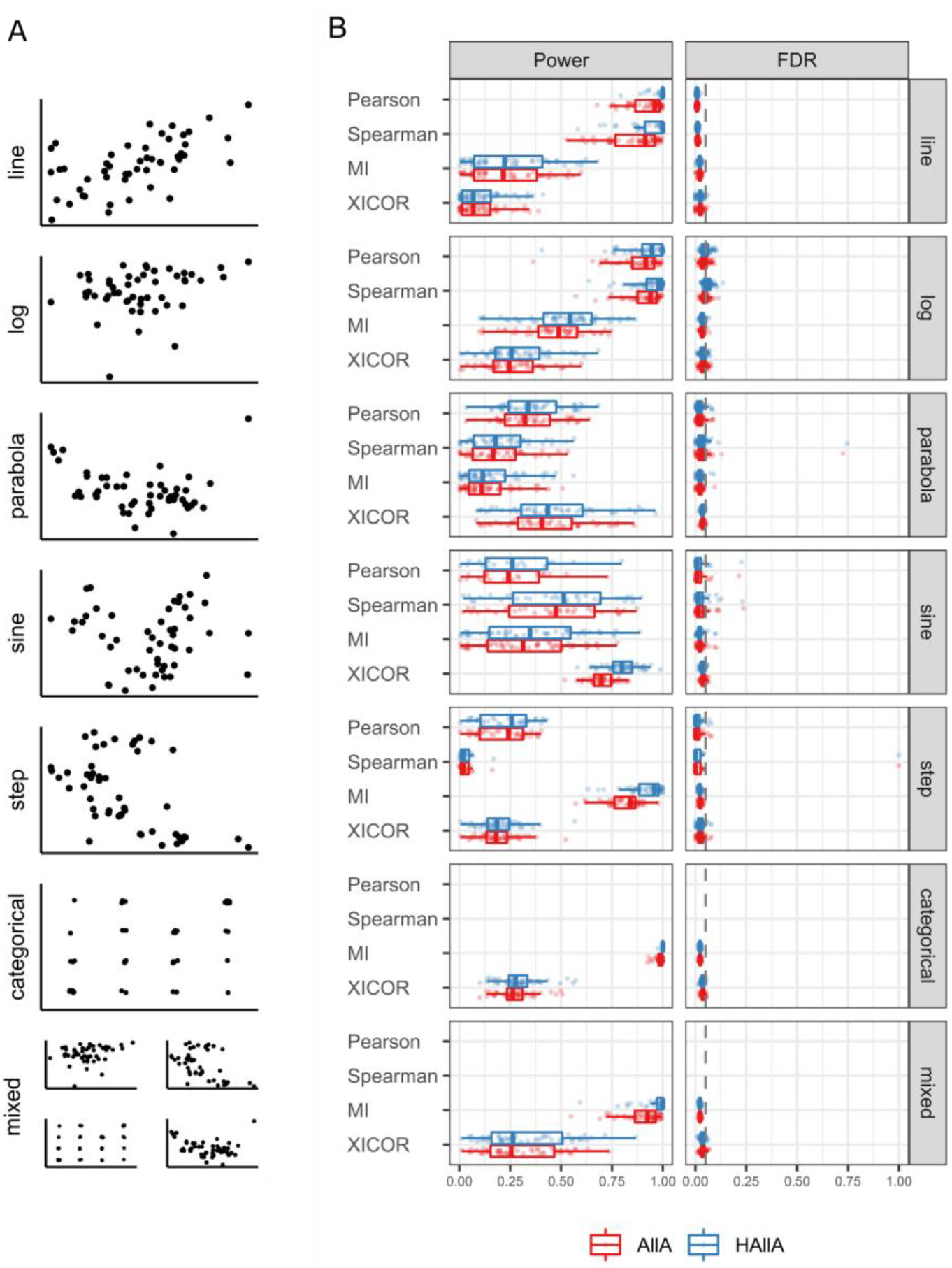
HAllA discovers block-structured associations while controlling false discovery rate. For a variety of feature linkage relationships, we simulated 50 independent paired datasets, each containing 200 features, 50 samples, and clusters of correlated features. We then evaluated the ability of hierarchical versus all-against-all testing to recover these associations using a variety of similarity metrics. Performance was evaluated by comparing power and false discovery rates. Our hierarchical all-against-all approach improved sensitivity relative to naive all-against-all approaches at a comparable false discovery rate. Similarity metrics that don’t accept categorical data have not been evaluated in the categorical or mixed association type. Other similarity metrics included in HAllA (dCor, NMI) were not applied in these simulations because their reliance on permutation tests made them too slow for simulations of this size (i.e. with many repeated iterations), although they are typically practical in individual real-world datasets.

We evaluated many different forms of feature association, including linear, quadratic, logarithmic, sinusoidal, stepwise, parabolic, and mixed (combined discrete and continuous) data. We compared HAllA and AllA across these association types using a variety of similarity measures, including XICOR, mutual information (MI), Spearman correlation, and Pearson correlation. Across datasets and similarity measures, HAllA consistently detected more built-in associations (had better average power by as much as 10%) than AllA while controlling FDR at the same pre-specified level (Fig 3B). Each similarity measure exhibited various strengths and weaknesses across evaluations depending on data type. As expected, for mixed and categorical data, MI is appropriate, and for monotonic associations in continuous data, Spearman correlation performs well. XICOR is applicable to both continuous and discrete outcomes and performs well on difficult nonlinear association types. However, it is rarely the most statistically powerful option, and its interpretation is limited to measuring the association of features in Y as a measurable function of features in X and not vice versa. A similar power analysis that used a fixed association structure with varying correlation strength led to similar conclusions (Fig 2 in S1 Appendix). Together these results show that the HAllA approach increases statistical power while maintaining the FDR across a wide variety of association structures under simulation.

### HAllA identifies novel fatty acid-xenobiotic metabolism associations in PPARα-deficient mice

PPARα is a nuclear receptor that regulates transcription of genes related to lipid metabolism in the liver [15]. These genes show high fatty acid catabolism rates, which can in turn affect hepatic fat storage and lipoprotein metabolism. We used HAllA to examine associations between 120 hepatic transcript levels and 21 liver lipid levels in a previously published dataset [16] (Fig 4). These data were originally collected from 40 wild type and peroxisome proliferator-activated receptor-α (PPARα)-deficient mice [15]. HAllA recovered 109 block associations comprising 225 pairwise associations at target FDR of 0.05 (chosen to match the previous study). HAllA’s results included all associations that were previously reported using canonical correlation analysis, including a key relationship between fatty acids and the xenobiotic metabolism genes Cyp3a11 and Car1(MGI:88268).

**Figure 4.**
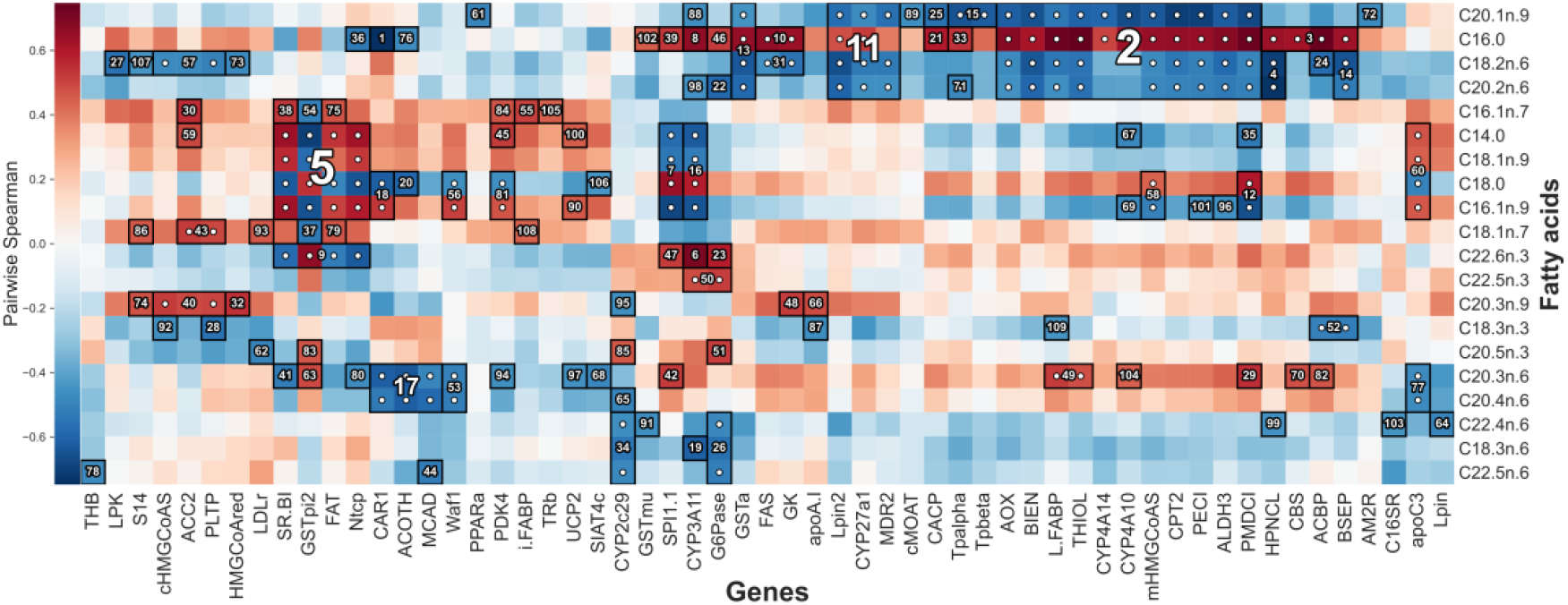
Association of fatty acids with host transcriptional activity in murine liver. We applied HAllA to paired data comprising 120 hepatic transcript levels and 21 liver lipid levels in a set of 40 previously profiled mice [15]. In this “HAllAgram” visualization of results, block associations are numbered in descending order of significance, with each numbered block corresponding to a group of co-expressed transcripts related to a group of co-occurring lipids. A white dot indicates marginal significance of a particular pair of features. A total of 109 block associations achieved significance at FDR 0.05, matching the previous study’s threshold based on canonical correlation [16] (detailed in S1 Table). HAllA’s associations were a strict superset of those found earlier by CCA. Spearman correlation was used as a similarity metric.

We further identified several novel associations, including a link between polyunsaturated fatty acids eicosatrienoic acid (C20:3n6) and arachidonic acid (C20:4n6) [17] with a group of transcripts including *Mcad* (*Acadm*, MGI:87867). This gene (C-4 to C-12 straight chain acyl-Coenzyme A dehydrogenase) encodes one of the main catalysts of the beta-oxidation process used for degradation of these fatty acids. Genes *Car1* (MGI:88268) and *Acot11* (MGI:1913736) (a carbonic anhydrase and lipid transfer protein, respectively [18-19]) fell in the same cluster with C20.3n.6 and C20.4n.6, which would suggest a trafficking and transport relationship between these genes and fatty acids.

### Associating microbes with metabolites in the infant gut microbiome

In a prior study, Kostic and colleagues examined the development of the human gut microbiome in a prospective, longitudinally sampled cohort of 33 Finnish and Estonian infants at high risk for type-1 diabetes [20]. Stool samples and clinical metadata (e.g. breastfeeding status, diet, and appearance of allergies) were collected monthly. Subjects’ stool samples were subsequently analyzed using 1) 16S rRNA amplicon sequencing (to profile gut microbiome composition) and 2) targeted mass spectrometry (to profile host and microbial metabolites). The dataset included 103 samples from 19 individuals, each with paired metabolomics and 16S rRNA gene sequencing data. We applied HAllA to identify associations between the residuals of microbial and metabolite abundances after correcting for longitudinal trends and subject specific random effects using a linear mixed effects model [21] (S1 Appendix).

HAllA recovered 44 microbial/metabolite cluster associations between 13 microbial genera and 44 metabolites using the same q < 0.05 threshold as in the original study (Fig 5A). These encompassed 57 pairwise associations, using Spearman correlation as the measure of pairwise feature similarity (as both data types are continuous). Using pairwise, all-against-all testing, 56 associations were significant at the same threshold.

**Figure 5.**
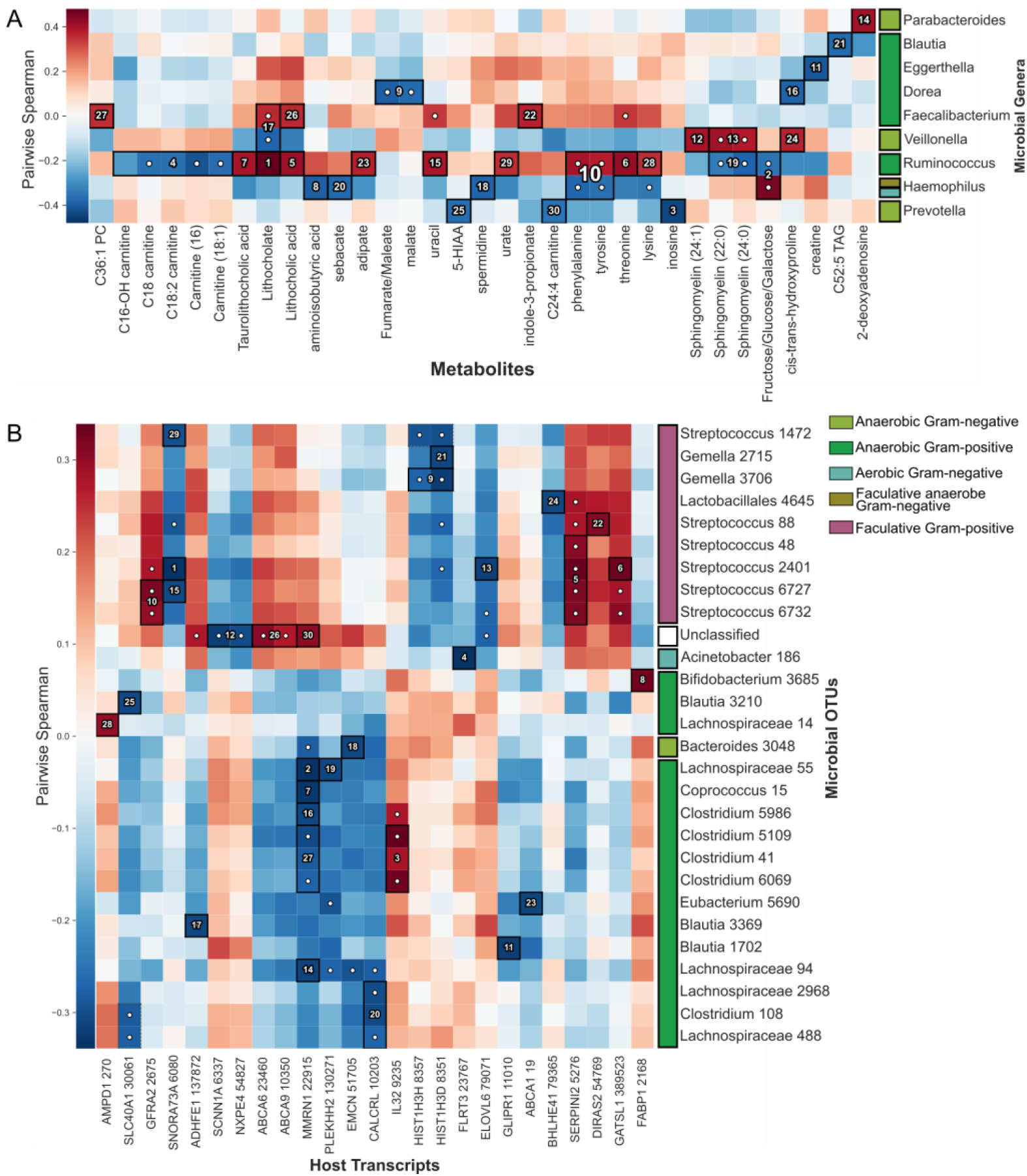
HAllAgram for block-wise associations. a) Using HAllA to associate multi-omic data for the analysis of metabolome-microbiome interactions. We used HAllA to associate paired stool metabolomic and 16S rRNA gene sequencing data from the DIABIMMUNE [20] cohort, in which infants were recruited at birth and sampled monthly for the first three years of life. The data comprise 104 samples and describes the abundance of 20 genera and 284 labeled metabolites. Here, we show the 30 strongest associations ranked by *p*-value (target FDR=0.05). **b) Relating host transcriptome and microbial taxa in IBD patients**. We applied HAllA to identify associations between the human gut microbiome and transcriptome in 204 patients receiving ileal pouch-anal anastomosis (IPAA) surgeries [23]. Block associations are numbered in descending order of significance based on best p-values in each block with each numbered block corresponding to a group of co-expressed transcripts related to a group of co-occurring microbial taxa (operational taxonomic units, OTUs).

Our results again replicate all significant associations from the previous study’s canonical correlation analysis (CCA), and most of the associations from the original pairwise association analysis of the previous paper. HAllA also found additional associations, including a novel association between *Prevotella* and inosine (Spearman coefficient = -0.439, FDR Q-value = 0.0053), which could be explained by a mechanism where increased levels of urotoxins in the body from inosine decreased the abundance of intolerant *Prevotella*. HAllA also reports novel associations between fecal bile acids lithocholate and lithocholic acid and genera *Faecalibacterium* and *Veillonella* (Spearman coefficients = 0.36, -0.39; Q-values = 0.026, 0.015, respectively). *Faecalibacterium* is Gram-positive anaerobic bacteria genera from order Clostridiales, while *Veillonella* are Gram-negative anaerobic cocci. Relationships between these genera and global bile acid levels (with matching correlation signs) has been previously indicated by several studies, particularly in cirrhosis [22]. These data thus demonstrate HAllA’s potential benefits relative to pairwise or omnibus (e.g. CCA) testing by simultaneously providing both greater interpretability and power.

### Associating the gut microbiome with host transcription in ulcerative colitis

We next applied HAllA to data combining 1) 16S rRNA amplicon sequencing of the human gut microbiome and 2) Affymetrix microarray screens of ileal RNA expression across 204 individuals in a cohort of ileal pouch-anal anastomosis (IPAA) patients [23]. In the original multivariate analysis of these data [24], microbial operational taxonomic unit (OTU) abundances were decomposed into principal components (PCs), and PCs accounting for up to 50% of the variance in the datasets were compared by all-against-all testing (an example of PC regression). While this approach enables well-powered comparisons of large numbers of features, the features are embedded as loadings in PCs, which complicates biological interpretation of the resulting associations.

HAllA identified 327 block associations in these microbial and gene expression data using an FDR threshold of 0.05 and a FNT of 0.1 (Fig 5B and S2 Table). Total relationships encompassed 125 OTUs, 187 transcripts, and the equivalent of 368 pairwise associations. The original study focused on the 9^th^ principal component (PC9) of the dataset due to its linking of a group of IL12/complement pathways to members of the microbiome, using an FDR threshold of 0.25. Of HAllA’s reported microbe-transcript associations when run with the same threshold, 20 genes were drawn from the 26 transcripts whose largest loading was in PC9. HAllA’s findings support a surprising result of the original study: although PC9 represented only 1% of the transcriptional variation in these samples, it captured most associations between transcription and the microbiome during pouchitis. These results also agree with a previous re-analysis of these data [25] assessing global covariation between gut microbial and transcriptional structure, which called out three pathways (interleukin-12, inflammatory, and inflammatory bowel disease genes) that overlap heavily with HAllA’s block results (e.g. 28 out of 51 tested genes in the KEGG TRP channel mediator pathway and 34 of 61 tested genes in the KEGG IBD pathway were significantly associated with microbial species).

Expanding on these previous associations, HAllA found a group of facultative anaerobes (mainly streptococci) to be positively associated with expression of the genes WDR49 and SERPINI2. WDR49 is a WD repeat-containing protein upregulated in alveolar macrophages, a cell type specifically responsible for nasopharyngeal pathogen uptake [26]. This association suggests this protein may also be involved in recognition of bacteria in the gut environment. Another novel association in HAllA’s results linked a group of *Bifidobacterium* OTUs with FABP1, a member of the long-chain fatty acid binding protein family involved both in lipid sensing and metabolic regulation of energy harvest [27]. This positive relationship has also been observed in mice [28]. Finally, during intestinal inflammation and bleeding, host-microbial iron competition is a limiting factor in subsets of microbial growth [29], which may be responsible for the significant negative association identified between the siderophore-rich genus *Blautia* and SLC40A1, a human intestinal epithelial iron ion transmembrane transporter [30].

### HAllA’s applicability to heterogeneous datasets

We finally applied HAllA to identify associations between mixed clinical metadata and RNA expression in the breast cancer cohort of the Cancer Genome Atlas (TCGA) [31] available from the LinkedOmics R package [32], focusing on highly expressed yet variable transcripts (Fig 3 in S1 Appendix). HAllA identified 483 significant (Q-value < 0.1) metadata-RNA associations within 261 blocks, including clusters of transcripts associated with tumor purity, PAM50 subtype, and ER Status. Notably, the transcripts occupying the block associated with PAM50 subtype include CA12, GABRP, NAT1, and TBC1D9, which have been previously proposed as predictor genes for breast cancer mortality, recurrence [33], and drug response [34]. Coupled with the results of the preceding applications, these results speak to the generality of HAllA’s association-discovery power across large, heterogeneous datasets.

In order to demonstrate the usefulness of alternative similarity measures like XICOR, we decided to look for non-linear functional relationships between RNA and protein expression in the breast cancer cohort of the Cancer Genome Atlas (TCGA) [31]. We applied HAllA to this data using both Spearman and XICOR as similarity measures, then examined the significant associations that came out with the latter but not the former. Among these we noticed three associations between RNA expression of transcription factor FOXC1 and protein expression of CCNE2, PIK3CA, and SRSF1 (FDR Q-value = 9.3×10^−7^, 3.9×10^−5^, 0.015, respectively) which showed compelling U-shaped relationships (Fig 6). When compared with PAM50 clinical subtypes, these relationships emerge as a result of two features of the originating tumors. First, the different PAM50 subtypes vary in average FOXC1 expression (i.e. average position on the x-axis). Secondly, the effect of FOXC1 on the expression of each protein appears to vary between the subtypes, with the opposite sign in the basal subtype. There are individually well-established links between subtype and FOXC1, CCNE2, and PIK3CA [35-37]. However, the varying relationship of each protein with FOXC1 by subtype has seemingly gone unnoticed in the literature, presumably due to the marginally non-linear shape of the overall relationship. While further study of the clinical importance of these relationships is warranted, these findings demonstrate the ease of well-powered, flexible, nonlinear association discovery with HAllA.

**Figure 6.**
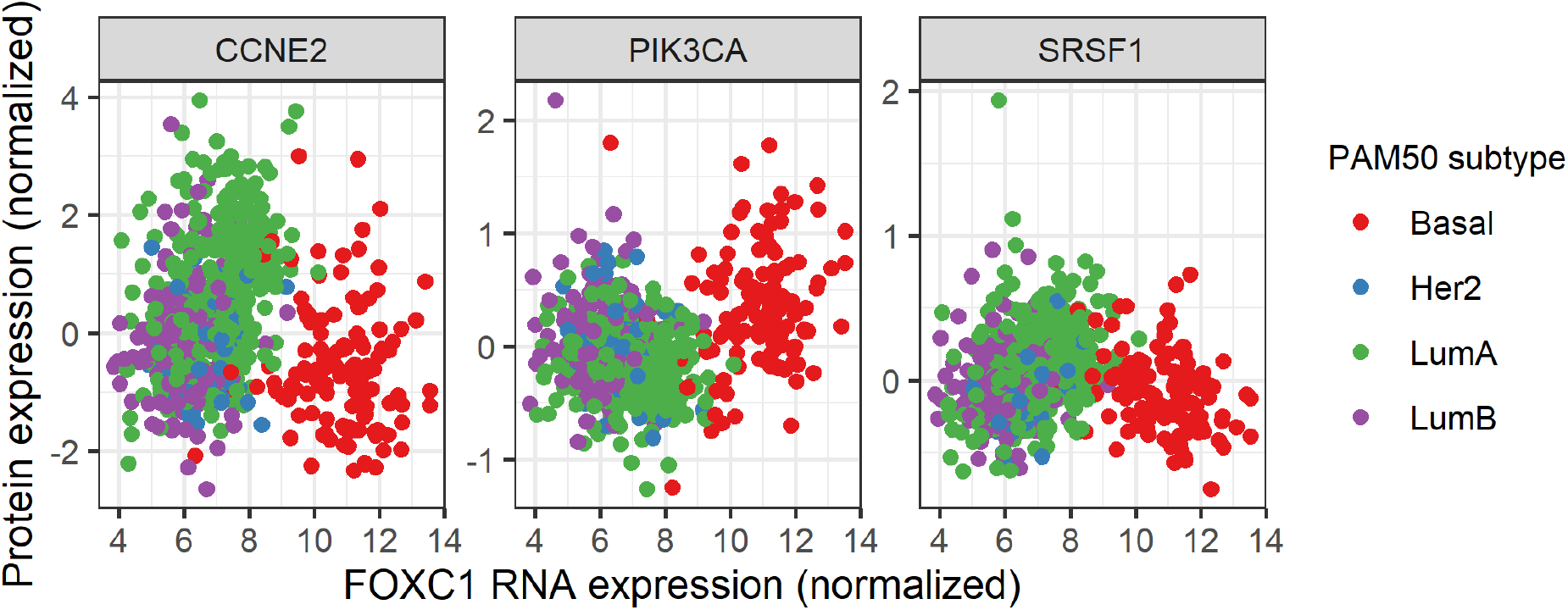
Non-linear relationships detected between RNA and protein expression in a breast cancer cohort. By using an association metric sensitive to nonlinear relationships (XICOR), HAllA detects U-shaped relationships between FOXC1 RNA expression and the protein expression of three genes. Overlaying the PAM50 subtype reveals that the U-shapes seem to emerge from a varying response to increased FOXC1 RNA expression by subtype. This effect seems to have gone unnoticed in the literature, thus demonstrating the ease with which HAllA can aid in the discovery of complicated relationships that might be missed otherwise.

## Discussion

In this work, we proposed and validated HAllA, a novel statistical method to find associations between multi-omic datasets. HAllA addresses several important methodological challenges in the analysis of high-dimensional datasets. It is applicable to data that are heterogeneous both within and between experiments, and it maintains statistical power using a novel hierarchical association testing and FDR control procedure. In this method, groups of correlated tests are modeled as blocks, ultimately reporting associations within blocks and between block representatives from multiple data types and experiments. This permits both great flexibility in the types of measurements to which it is applied and ease of interpretation of the resulting significant associations.

Class prediction approaches are commonly used to model relationships between high-dimensional datasets with variables measured using shared observational units. For example, Partial Least Squares [38] and its close relative Canonical Correlation Analysis [39] identify latent variables in one dataset that are maximally correlated to latent variables in the other dataset. These methods, and robust and penalized varieties [40-41], can identify blocks of variables that are correlated within one dataset and in turn with another block of correlated variables in another dataset. They do not, however, control for family-wise error or FDR, and so are most suitable for prediction or exploratory, visual, and descriptive analysis. With these methods, inference on the existence of associations between the variables of two datasets against null hypotheses of independence still relies on univariate hypothesis tests (and possibly dimension reduction or clustering) and is performed subsequently in a separate step. The FDR for the potentially large number of tests can be controlled by the Benjamini and Hochberg method [42], which has been adapted for dependent tests [43] and hierarchically organized tests [44] that are continued until non-significance. The approach described here thus aims to combine the best features of these different existing approaches, yielding clustering of potentially heterogeneous variable types within each dataset with hierarchical testing and control of FDR.

While these approaches are frequentist, Bayesian models are also used to improve power and share information among feature blocks [45-48]. While such methods are extremely powerful within their target domains, they are typically intended for incorporation of specific prior knowledge, such as graph structure [44, 49], phylogeny [50], or pathway-based functional roles [51]. They can also be computationally expensive in cases where many or long simulation chains are required for convergence [52]. HAllA’s nonparametric frequentist approach will likely result in reduced power relative to such models within the domains for which they are designed, but with substantially reduced computational cost and without the need to specify model relationships and priors in each new application domain. Like most statistical tradeoffs, HAllA’s generality as a tool for association discovery thus comes at a cost in specific circumstances where it is desirable to instead utilize prior knowledge and known data structure.

A limitation of the current method is that it can only look for associations between two datasets at a time. While the method can be applied to multiple pairs of joint datasets manually, this becomes combinatorially prohibitive in particularly thorough studies where a large number of high-dimensional data types are available (e.g. studies which collect genetics, gene expression, epigenetics, microbial profiles, metabolites, and metadata from each sample). In circumstances such as these, repeated application of HAllA across each pair of datasets would no longer properly control FDR. A potential extension would be to incorporate multivariate testing directly as an association measure, e.g. block PERMANOVA [53-54] or Procrustes analysis [55], to lower the combinatorial burden by performing inference on sets of features rather than individual feature pairs. Second, the model does not share information between blocks, as would be the case in a fully multivariate test [53] or a hierarchical Bayesian model [48]. Cases in which data do include such multi-layered nonindependence structure may indeed be better handled in a Bayesian framework. Finally, and relatedly, it is not straightforward to incorporate any type of prior knowledge into the HAllA framework, again because of HAllA’s intention for wide applicability. Pre-filtering can be used, as in several of our own examples, but this can be either beneficial or detrimental depending on context [56-57].

Future work could also provide several refinements to the method, in addition to addressing these limitations. Currently, for example, known but undesirable confounders must be separately regressed out prior to using HAllA, and the method run on the resulting residuals instead of raw data. Integrating such covariate adjustment would be possible in future versions of the method’s implementation. Perhaps most importantly, it may be possible to place tighter theoretical bounds on the block-wise and global FDR control beyond what is provided by HAllA’s adaptation of the Benjamini-Hochberg [42] and Benjamini-Yekutieli methods [58]. This would also suggest a theoretical framework within which to characterize the amount and types of non-independence best handled by hierarchical block association testing. Ultimately, tradeoffs must be made between power and generality [59]. However, we aim for HAllA to provide a happy medium, capable of serving as an easy-to-use first pass analysis in a wide range of multi-omics data types.

## Acknowledgements

We thank Alex Kostic, Tommi Vatanen, and Vincent Carey for assistance obtaining and curating datasets for the applications section; Hera Vlamakis, Hector Corrada Bravo, William Shannon, A. Brantley Hall, Himel Mallick, Siyuan Ma, and Susan Holmes for helpful discussions, suggestions, and feedback. This study was supported by Army Research Office grant W911NF-11-1-0429, NSF DBI-1053486, and NIH U54DE023798 to Curtis Huttenhower.

## Author Contributions

G.R., E.F., L.W., and C.H. conceived the method; G.R., K.S., and A.G. implemented the software; G.R., K.S., A.G., and L.M. tested and packaged the software. G.R., A.G., and E.F. evaluated the performance; G.R., G.W., A.G., and L.M. provide online documents and software. G.R., J.L.-P., Y.M., A.G., and X.M. prepared synthetic data and applications. G.R., E.F., A.G. and C.H. wrote the manuscript. All authors discussed the results and commented on the paper.

## Supporting information

### S1 Appendix. Supplementary methods and evaluation

**S1 Table. HAllA results on data from PPARα-deficient mice**

Significant HAllA results with FDR threshold q = 0.05 for fatty acid-transcript associations in PPARα-deficient mice [15].

**S2 Table. HAllA results on microbe-gene relationships**

Significant HAllA results with FDR threshold q = 0.1, Spearman correlation as similarity metric, and medoid as the decomposition method for microbial and gene expression profiling data [23]. Reported associations encompassed 427 OTUs, 1,991 transcripts, and the equivalent of 8,382 pairwise associations.

**S3 Table. HAllA results microbe-metabolite relationships**

Significant HAllA results with FDR threshold q = 0.25, Spearman correlation as the similarity metric, and medoid as decomposition method, for the DiabImmune cohort data from [21]. These include 20 microbial genera and 284 metabolites of 103 samples.

